# Crystal structure of human endothelin ETb receptor in complex with peptide inverse agonist IRL2500

**DOI:** 10.1101/460410

**Authors:** Chisae Nagiri, Wataru Shihoya, Asuka Inoue, Francois Marie Ngako Kadji, Junken Aoki, Osamu Nureki

## Abstract

Endothelin receptors (ET **A** and ET **B**) are G-protein coupled receptors activated by endothelin-1 and are involved in blood pressure regulation. IRL2500 is a peptide-mimetic of the C-terminal tripeptide of endothelin-1, and has been characterized as a potent ET **B**-selective antagonist, which has preventive effects against brain edema. Here, we report the crystal structure of the human ET **B** receptor in complex with IRL2500 at 2.7 A-resolution. The structure revealed the different binding modes between IRL2500 and ET-1, and provides structural insights into its ET **B**-selectivity. Notably, the biphenyl group of IRL2500 penetrates into the transmembrane core proximal to D2.50, stabilizing the inactive conformation. Using the newly-established constitutively active mutant, we clearly demonstrate that IRL2500 functions as an inverse agonist for the ET **B** receptor. The current findings will expand the chemical space of ETR antagonists and facilitate the design of inverse agonists for other class A GPCRs.

## Introduction

Endothelin receptors (ETR) are G-protein coupled receptors activated by vaso active peptide, endothelins **^1^**. Two endothelin receptor subtypes (ET **A** and ET **B**) are widely expressed in the vascular endothelium, brain, and other circulatory organs **^2, 3^**. Endothelin-1 (ET-1) activates the endothelin receptors (ETRs) with sub-nanomolar affinities. The activation of the ET **A** receptor leads to potent and long-lasting vasoconstriction, whereas that of the ET **B** receptor induces nitric oxide-mediated vasorelaxation. Therefore, the up-regulation of ET-1 is significantly related to circulatory-system diseases, including pulmonary arterial hypertension (PAH) **^4-7^**. Moreover, the autocrine and paracrine signaling functions of ET-1 through the ET **A** receptor play a critical role in tumor growth and survival **^8^**. Therefore, ETR antagonists have been developed for the treatment of circulatory-system diseases and cancers **^6, 7^**. Bosentan is the first orally-active ETR antagonist **^9, 10^**, and is used to treat PAH. The ET **B** receptor is the prominent ET receptor subtype in the brain, with high expression levels in astrocytes **^11^**. Stimulation of the ET **B** receptor modulates astrocytic responses, indicating its important roles in regulating astrocytic functions **^12^**. The up-regulation of the astrocytic ET **B** receptor by ET-1 increases the vascular permeability and reduces the AQP4 levels, thereby aggravating vasogenic brain edema **^11^**. The application of ET **B**-selective antagonists may provide preventive effects against brain edema in the acute phase of brain insults **^13-16^**.

To date, most endothelin receptor antagonists have been developed based on bosentan **^17, 18^**. The ETR antagonists that have been developed till now are mostly N-heterocyclic sulfonamides with similar structures and molecular weights, and nonsulfonamide antagonists (atrasentan, ambrisentan, darusentan, and enrasentan) still retain high similarities with each other and with the sulfonamides **^7^**. Therefore, the ETR agents are chemically very similar, and expanded chemical space should be exploited. IRL2500 is a peptide ETR antagonist developed based on the partial structure of ET-1 **^19^**, rather than bosentan. IRL2500 has been characterized as an ET **B**-selective antagonist with an IC **50**value of 1.2 nM **^20^**, which shows higher affinity than that of bosentan. In an animal model, the intracerebroventricular administration of IRL2500 attenuated cold injury-mediated brain edema and disruption of the blood-brain barrier, indicating the neuroprotective effect of IRL2500 **^14, 15^**. An understanding of the IRL2500 binding mode would facilitate the expansion of the chemical space of ET agents.

We previously reported the crystal structures of the ET **B** receptor bound to ET-1 **^21^**and bosentan **^22^**; however, both the binding mode and ET **B**-selectivity of IRL2500 remained to be elucidated. Here, we present the crystal structure of the ET **B** receptor in complex with IRL2500. This structure revealed the unique binding mode of IRL2500, which differs from those of ET-1 and bosentan. Structure-guided functional analyses clearly demonstrate that IRL2500 functions as an inverse agonist for the ET **B** receptor, and thus will provide the basis for design of inverse agonists for other class A GPCRs.

## Results

### Overall structure

For crystallization, we used the previously established, thermostabilized ET **B** receptor (ET **B**-Y4) **^22, 23^**. The IC **50**value of IRL2500 for ET **B**-Y4 was similar to that for the wild type receptor in the TGFa shedding assay **^24^**(Fig. 1), suggesting that the themostabilizing mutations minimally affect the IRL2500 binding. In contrast, the IC **50**value of IRL2500 for the ET **A** receptor is over 3 (Fig. 1), indicating that IRL2500 has over 100-fold ET **B**-selectivity, consistent with the previous pharmacological analysis **^20^**. To facilitate crystallization, we replaced the third intracellular loop (ICL3) of the receptor with minimal T4 Lysozyme **^25^**(ET **B**-Y4-mT4L). Using ***in meso*** crystallization **^26^**, we obtained crystals of ET **B**-Y4-mT4L in complex with IRL2500 (Supplementary Fig. 1a, b). In total, 58 datasets were collected and merged by the data processing system KAMO **^27^**. Eventually, we determined the ET **B** structure in complex with IRL2500 at 2.7 A resolution, by molecular replacement using the antagonist-bound ET **B** structure (PDB code: 5X93) (Table 1).

**Table 1.**
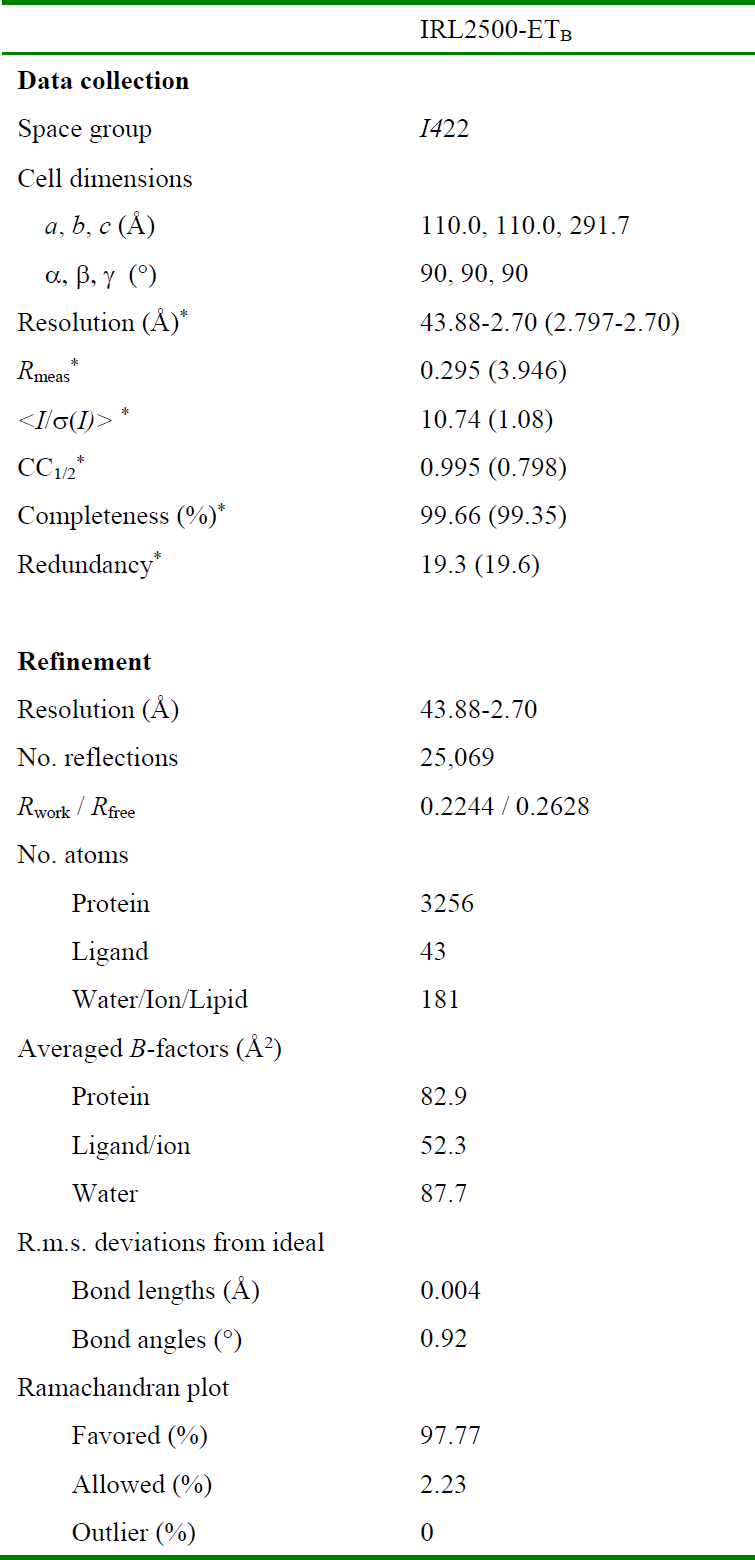

**Fig S1.**
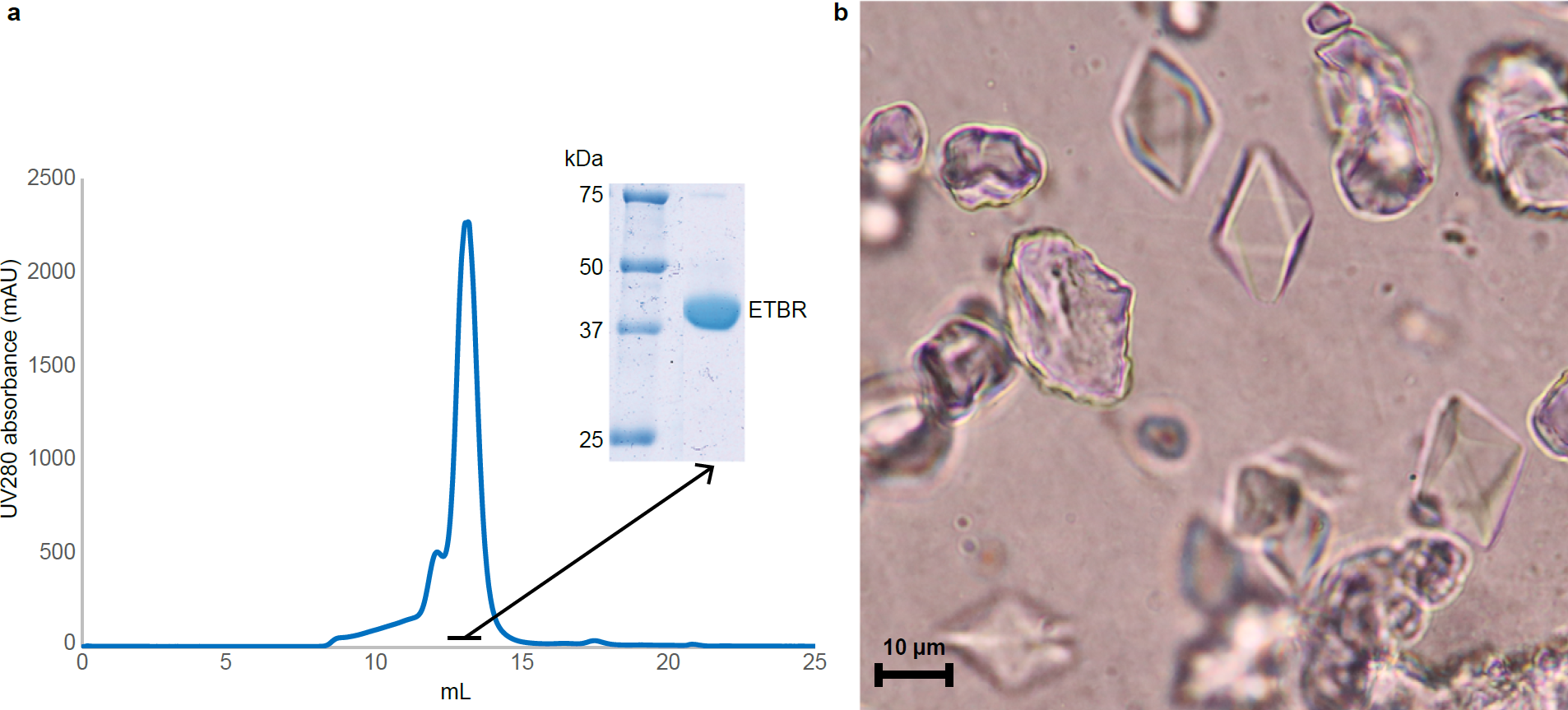
Supplementary Fig. 1. Crystallization. **a,**Gel filtration chromatogram and SDS-PAGE of the purified IRL2500-bound ET **B** receptor. **b**, Crystals of the IRL2500-bound ET **B** receptor.

**Fig.1.**
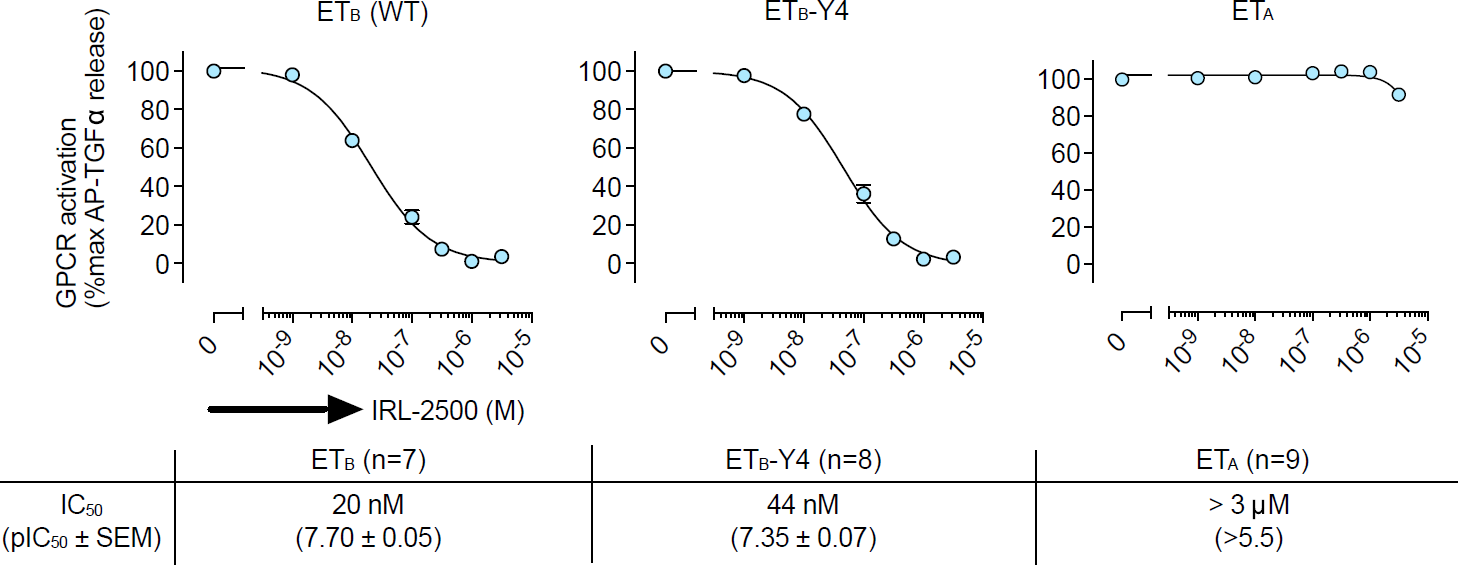
Inhibition of ET-1 binding by IRL2500. Effect of IRL2500 on the ET-1 (0.2 nM)-induced release of AP-TGFa in HEK293 cells expressing the endothelin receptors. For each experiment, the AP-TGFa release response in the absence of IRL2500 is set at 100%. Data are displayed as means ± s.e.m. (standard error of the mean) from seven to nine independent experiments, with each performed in triplicate.

The overall structure consists of the canonical 7 transmembrane helices (TM), the amphipathic helix 8 at the C-terminus (H8), and two antiparallel β-strands in the extracellular loop 2 (ECL2), as in the previously determined ET **B** structures (Fig. 2a). The IRL2500-bound structure is similar to the bosentan-bound structure, rather than the ET-1-bound structure (R.M.S.D. values for Cα atoms=1.34 and 1.95 A, respectively), reflecting the inactive conformation. We observed a remarkable difference in the conformation of ECL2. The β strands are opened up by 9 Å, as compared with those in the ligand-free structure (Fig. 2b and Supplementary Fig. 2a), and are the widest among the peptide-activated class A GPCRs (Supplementary Fig. 2b). This structural feature indicates the innate flexibility of ECL2, to capture the large peptide ligand endothelin, in the inactive conformation of the ETB receptor.

**Fig S2.**
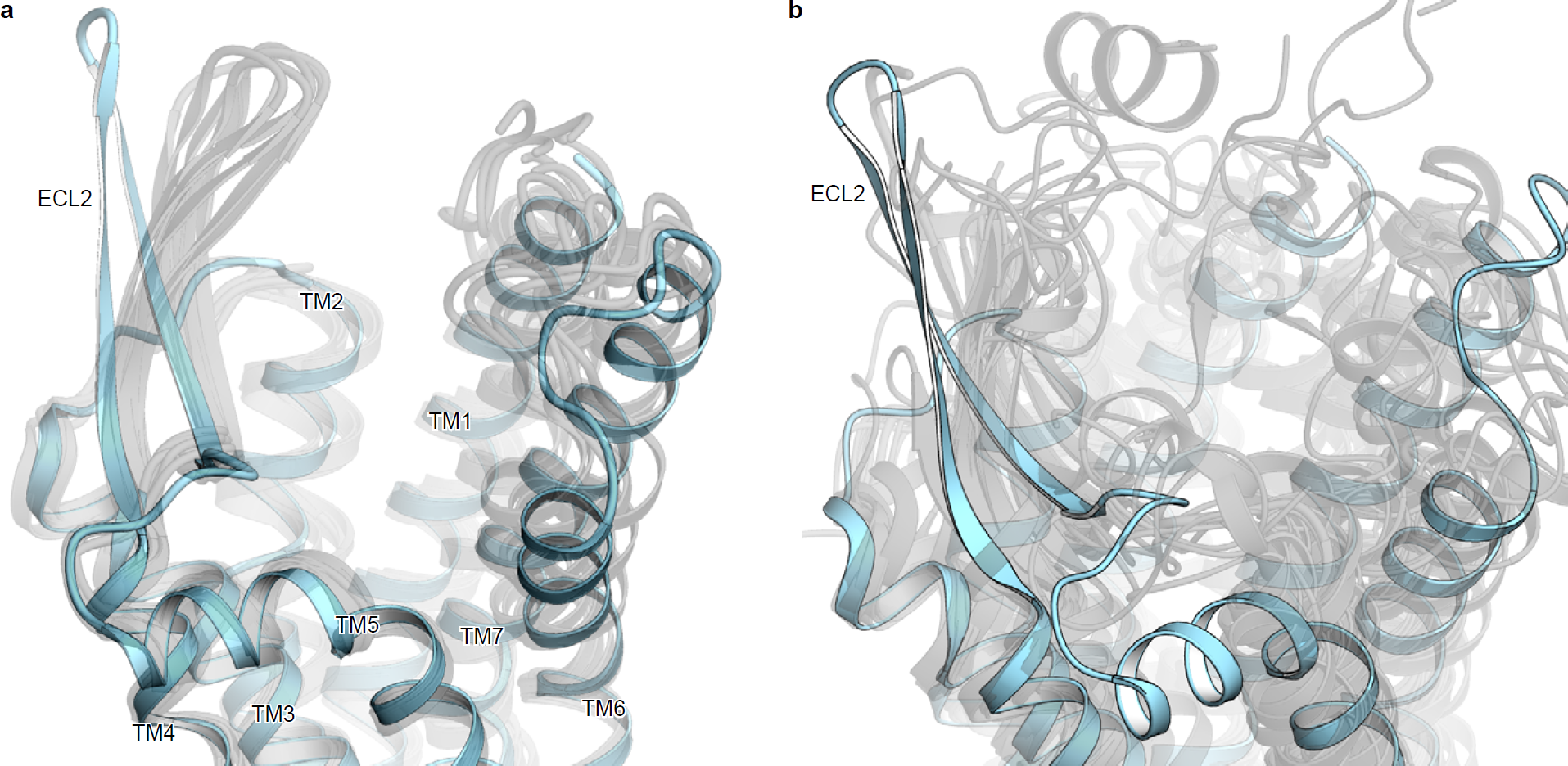
Supplementary Fig. 2. Comparison of the ECL2 structure with those of other peptide-activated GPCRs. **a**, Superimposition of the ET **B** structures determined to date. The IRL2500-bound and other structures are coloured sky blue and gray, respectively. **b**, Superimposition of the IRL2500-bound ET **B** structure with other peptide-activated class A GPCRs.

**Fig.2.**
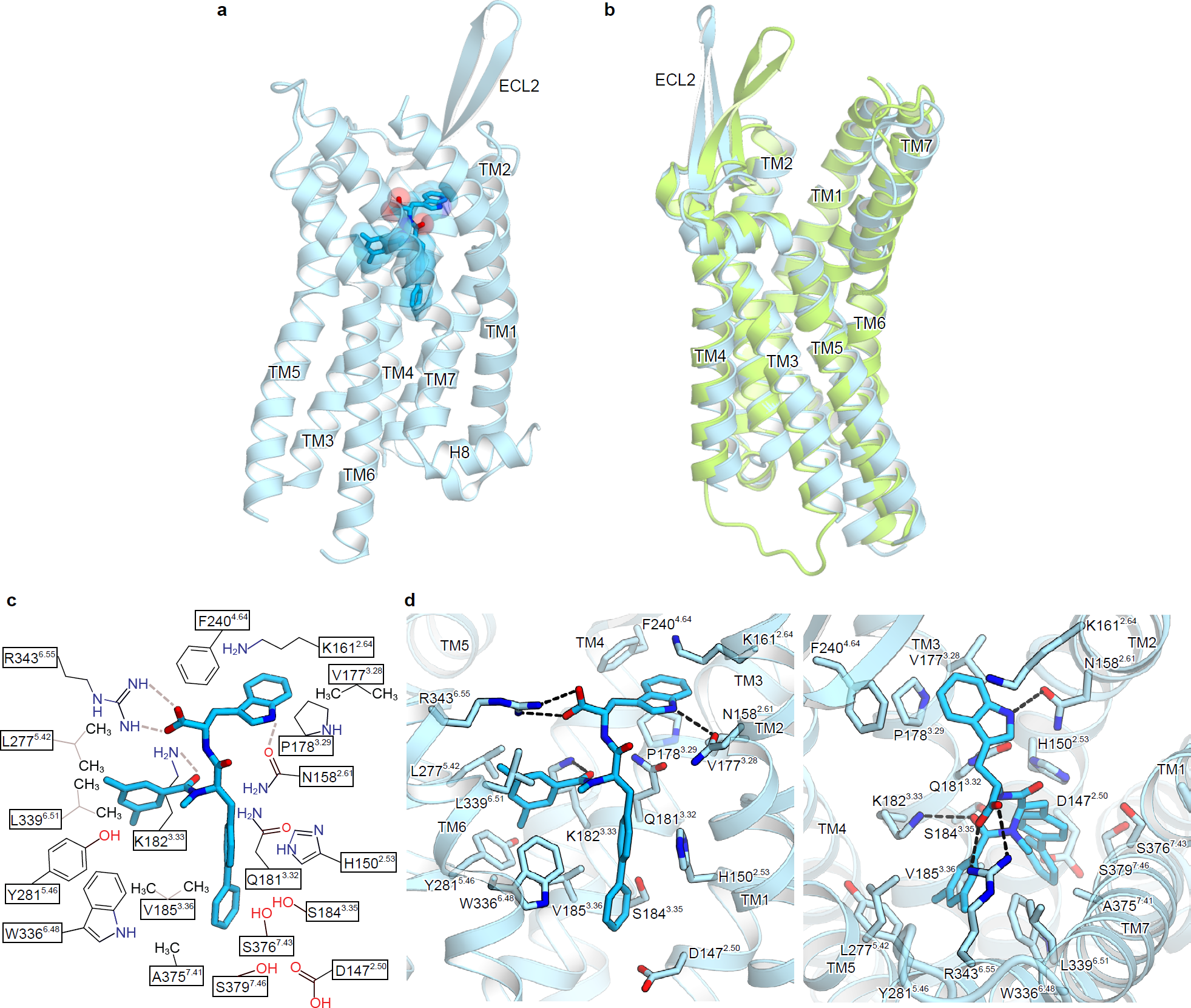
ETb structure in complex with IRL2500. **a**, The overall structure of the IRL2500-bound ET **B** receptor. The receptor is shown as a sky blue ribbon model. IRL2500 is shown as a deep sky blue stick with a transparent surface model. **b**, Superimposition of the IRL2500-bound and ligand-free ET **B** structures (PDB code: 5GLI), colored sky blue and light green, respectively. **c**, Schematic representation of the interactions between ET **B** and IRL2500 within 4.5 A. The dashed lines show hydrogen bonds. **d,**Binding pocket for IRL2500, viewed from the extracellular side (over) and the membrane plane (lower). The receptor is shown in a sky blue ribbon representation. IRL2500 and receptor residues involved in ligand binding are shown as sticks, coloured deep sky blue and sky blue, respectively. The dashed lines show hydrogen bonds.

### IRL2500 binding site

We first describe the IRL2500 binding mode. IRL2500 consists of a tryptophan, a 3,5-dimethylbenzoyl group, and a biphenyl group **^19^**, which are connected by two peptide bonds (Fig. 2c). IRL2500 binds to the transmembrane binding cleft exposed to the extracellular side, with a clear electron density in the ***Fo-Fc*** omit map (Supplementary Fig. 3a, b). The carboxylate group of the tryptophan moiety in IRL2500 forms salt bridges with K182 **^3.33^** and R343 **^6.55^**(superscripts indicate Ballesteros-Weinstein numbers **^28^**) (Fig. 2c, d). The tryptophan side chain of IRL2500 hydrogen bonds with the carbonyl group of the N158 **^2.61^**side chain, and forms extensive van der Waals interactions with N158 **^261^**, K161 **^2.64^**, V17 7 **^3.28^**, P178 **^3.29^**, and F240 **^4.64^**(Fig. 2d). The dimethyl phenyl group of IRL2500 forms van der Waals interactions with the hydrophobic pocket, and is surrounded by V185 **^3.36^**, L27 7 **^5, 42^**, Y28 1 **^5, 46^**, W3 3 6 **^6, 48^**, L339 **^6.51^**, and H340 **^6.52^**The biphenyl group penetrates deeply into the receptor core proximal to D147 **^2, 50^**, and forms van der Waals interactions with D147 **^2, 50^**, H15 0 **^2, 53^**, W336 **^6.48^**, and S376 **^7.43^**. Overall, the carboxylate of IRL2500 is specifically recognized by the positively charged residues of the ET **B**receptor, and the other moieties fill the space within the transmembrane binding pocket.

**Fig S3.**
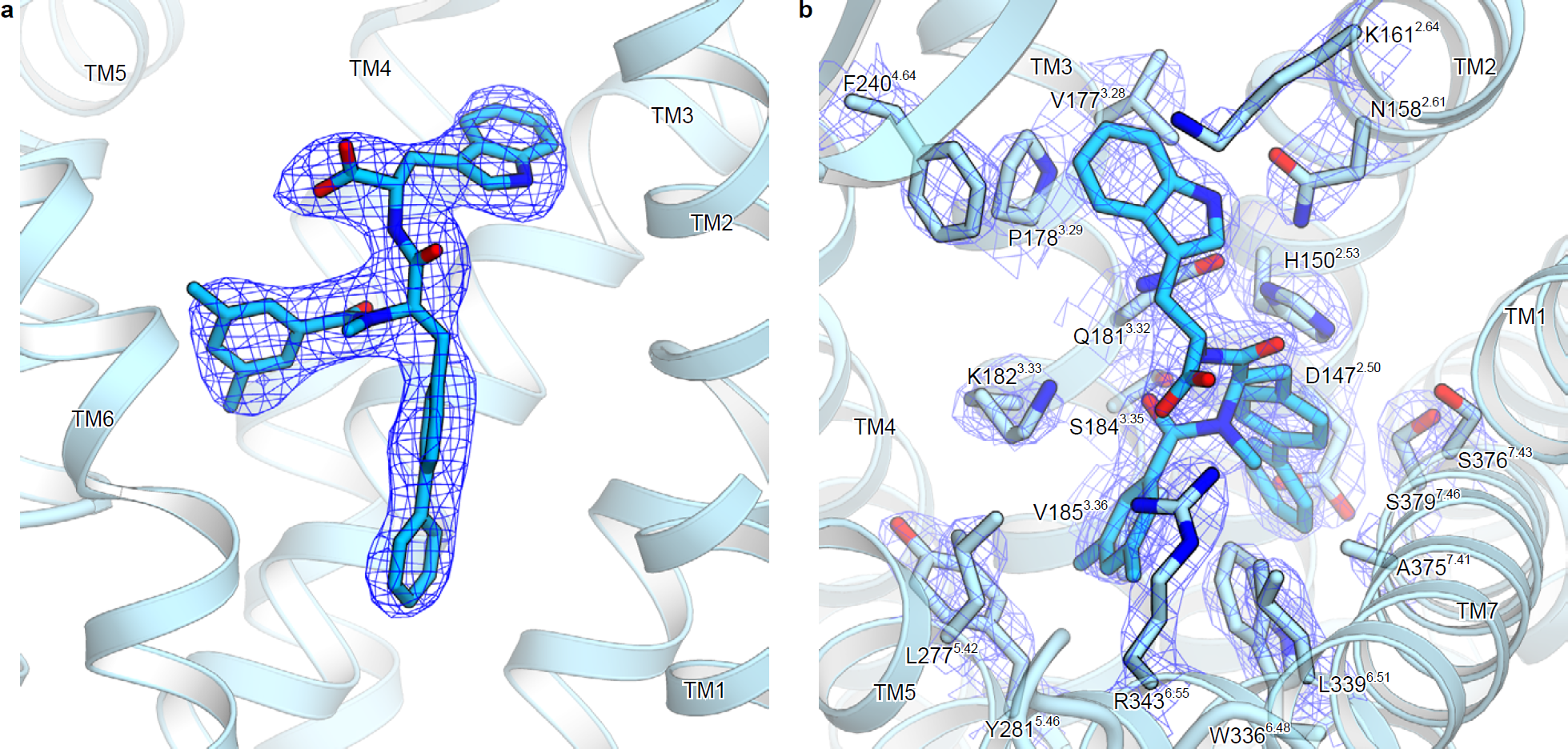
Supplementary Fig. 3. Electron density. **a**, ***F****o* ***-F****c* omit maps for IRL2500, contoured at 3.0o. **b**, ***2F****o* ***-F****c* map around the IRL2500 binding site, contoured at 2.5o.

To elucidate the structural basis for the ET **B**-selectivity of IRL2500, we compared the residues constituting the IRL2500 binding site between the ET **B** and ET **A** receptors (Fig. 3a and Supplementary Fig. 4). Although most of the residues are conserved, three residues are replaced with bulkier residues in the ET **A** receptor (H150Y, V177F, and S376T). These replacements may cause steric clashes with the aromatic groups of IRL2500 and reduce its affinity. To investigate this hypothesis, we measured the IC **50**values of IRL2500 for the H150Y, V177F, and S346T ET **B** receptor mutants. These mutants showed similar responses for ET-1 in the TGFα shedding assay (Supplementary Fig. 5), and only V177F showed a 4-fold higher IC **50**value with IRL2500 (Fig. 3b). These data suggest that the V177F mutation in the ET **A**receptor sterically clashes with the tryptophan moiety of IRL2500 and reduces its affinity, thus partially accounting for the ET **B**-selectivity of IRL2500. This is consistent with the previous study, in which the replacement of the tryptophan moiety with the smaller valine residue in IRL2500 weakened its ET **B**-selectivity **^29^**.

**Fig S4.**
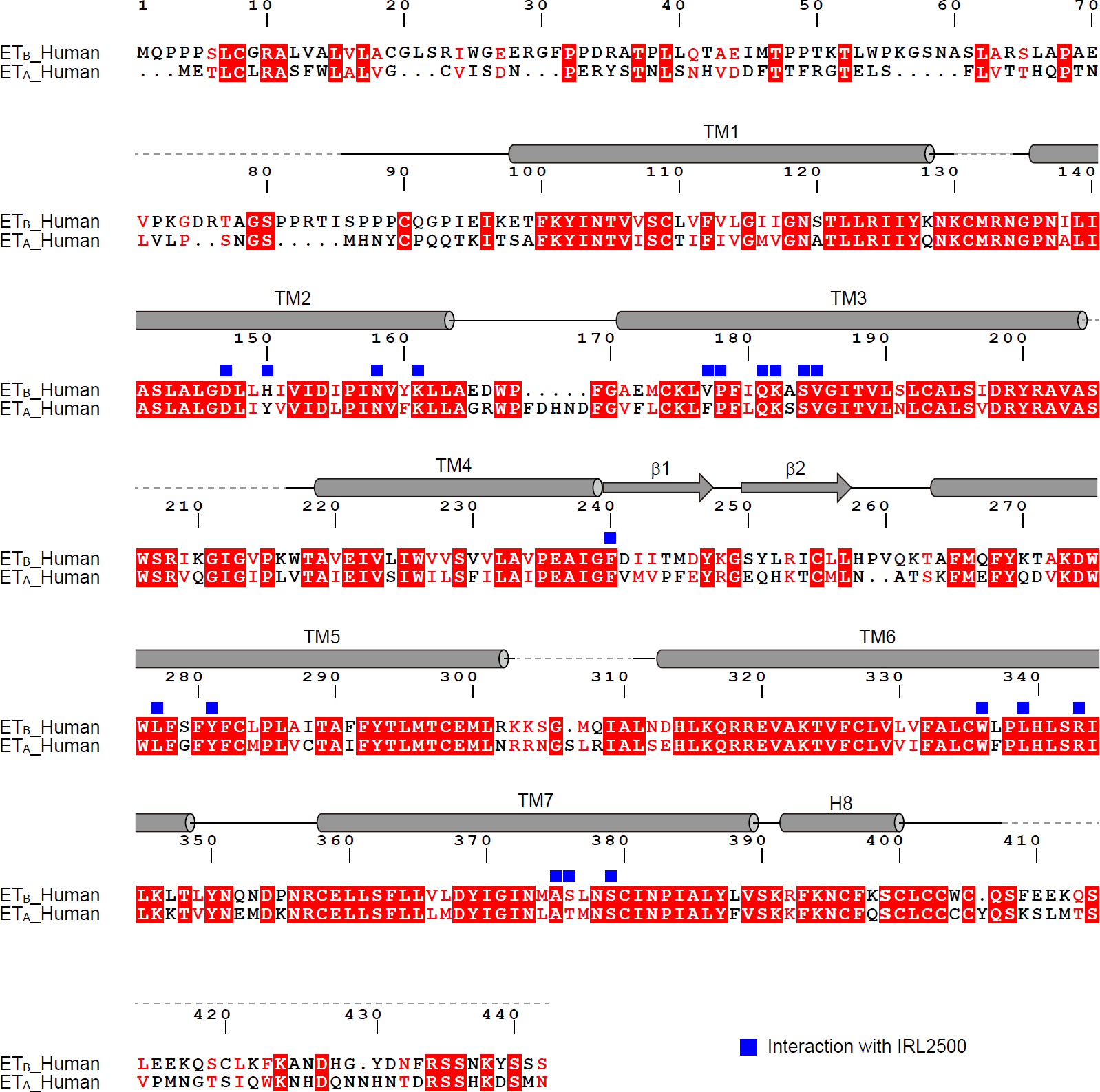
Supplementary Fig. 4. Alignment of the human ETa and ETb receptors. Alignment of the amino acid sequences of the human ET **B** receptor (UniProt ID: P24530) and the human ET **A** receptor (P25101). Secondary structure elements for a-helices and P-strands are indicated by cylinders and arrows, respectively. Conservation of the residues between ET **A** and ET **B** is indicated as follows: red panels for completely conserved; red letters for partially conserved; and black letters for not conserved. The residues involved in IRL2500 binding are indicated with squares.

**Fig S5.**
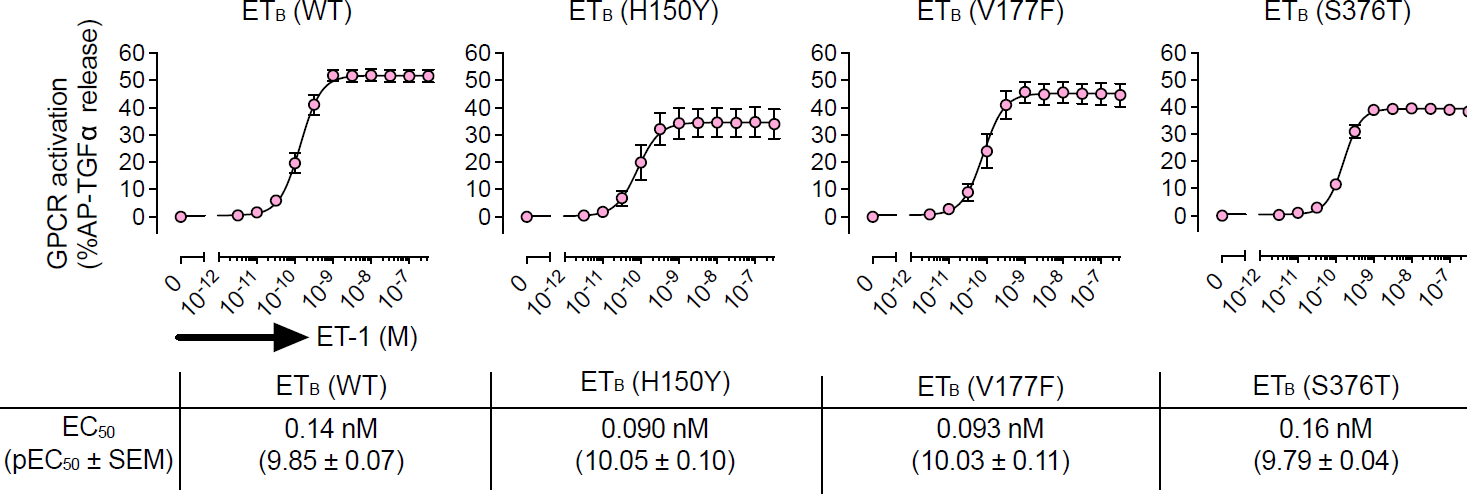
Supplementary Fig. 5. ET-1 responses of mutant receptors. Concentration response-curves of AP-TGFa release in the ET-1 treatment of HEK293 cells expressing the endothelin receptors. Symbols and error bars are means and s.e.m. (standard error of the mean) of four or six independent experiments, each performed in triplicate.

**Fig.3.**
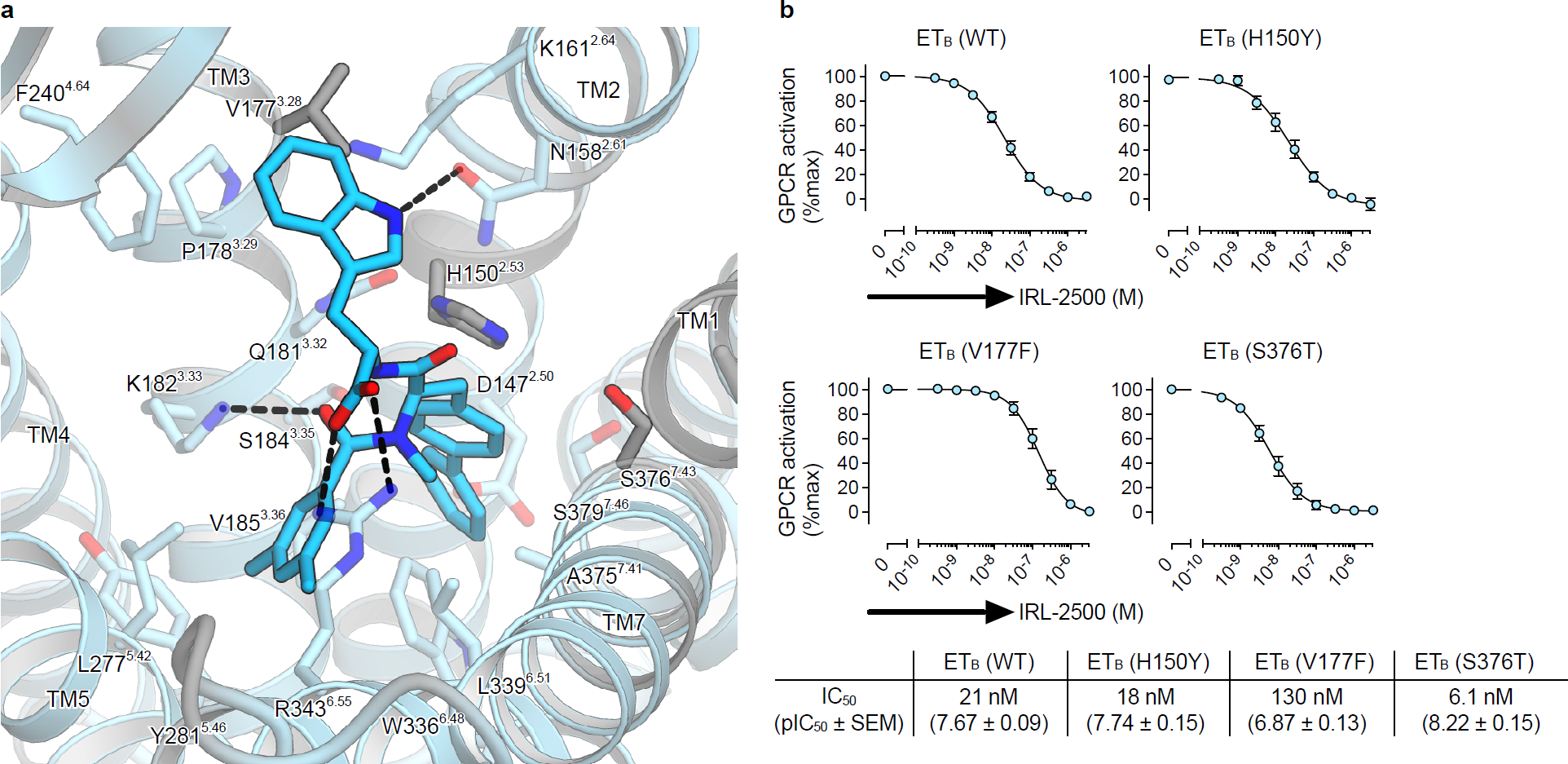
Conservation of the IRL2500 binding site. **a**, Sequence conservation of the IRL2500 binding site between ET **A** and ET **B**, mapped onto the IRL2500-bound structure. Conserved and non-conserved residues are coloured sky blue and gray, respectively. The receptor residues involved in IRL2500 binding are shown as sticks. The dashed lines show hydrogen bonds. **b**, Effect of IRL2500 on the ET-1 (0.2 nM)-induced release of AP-TGFa in HEK293 cells expressing the mutant ET **B** receptors. For each experiment, the AP-TGFa release response in the absence of IRL2500 is set at 100%. Data are displayed as means±s.e.m. (standard error of the mean) from four to six independent experiments, with each performed in triplicate.

### Comparison of the binding modes of IRL2500, ET-1, and bosentan

IRL2500 is designed to mimic the Y13, F14, I19, I20, and W21 residues in ET-1, which play critical roles in ligand binding to the ET **B** receptor **^19^**. The tryptophan and dimethyl phenyl group of IRL2500 seem to be equivalent to W21 and I20 in ET-1, respectively, while the biphenyl group of IRL2500 seems to be equivalent to F14 and I19 of ET-1. However, a comparison between IRL2500 and ET-1 binding revealed an unexpected difference in their binding interactions (Fig. 4a). The carboxylate of the tryptophan in IRL2500 superimposes well with that of W21 in ET-1, and is coordinated by similar positively charged residues. The tryptophan moiety and dimethyl phenyl group of IRL2500 superimpose well with I20 and W21 of ET-1, respectively. In contrast, the biphenyl group of IRL2500 penetrates into the receptor core, in an opposite manner to the F14 and I19 of ET-1. Overall, the electrostatic interactions between the carboxylates and the positively charged residues are conserved in IRL2500 and ET-1 binding, but the other moieties form totally distinct interactions with the receptor. The volume of the ligand binding pocket in the ligand-free structure is large, thereby allowing the aromatic moieties of IRL2500 to flip.

**Fig.4.**
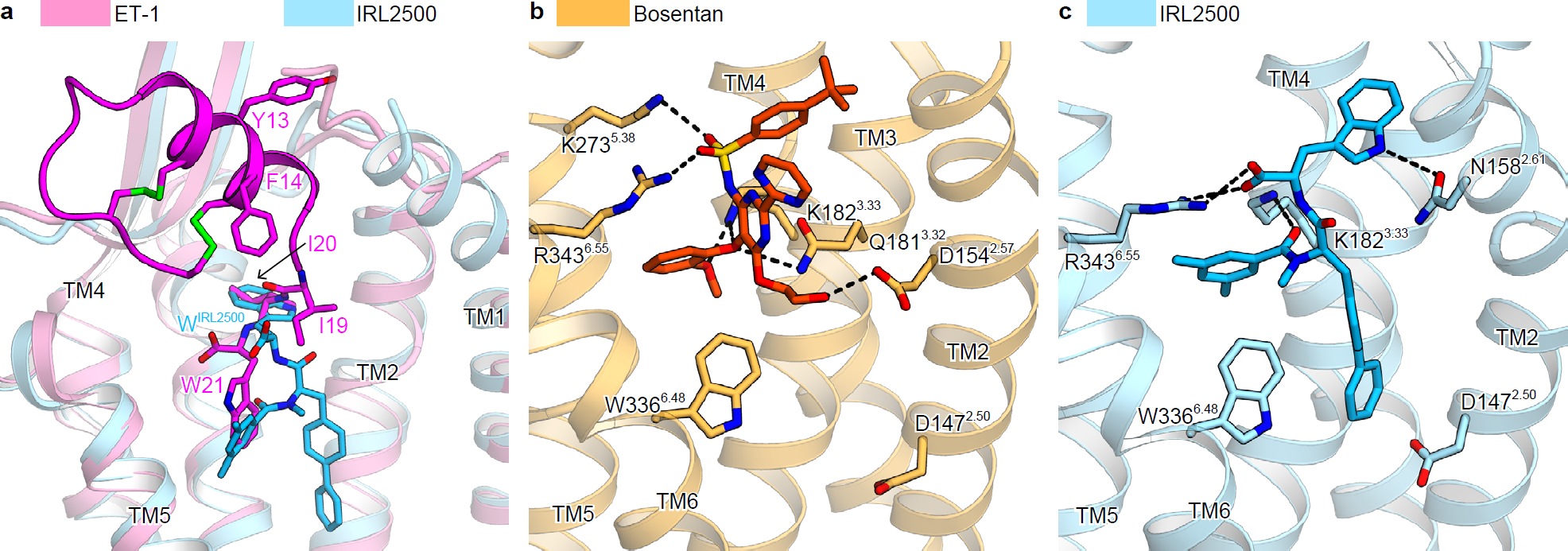
Comparison of binding modes of IRL2500, ET-1, and bosentan. **a,**Superimposition of the IRL2500-and ET-1-bound ET **B** receptors (PDB code: 5GLH). The ET-1-and IRL2500-bound receptors are shown as pink and sky blue ribbons, respectively. IRL2500 is shown as a stick model. ET-1 is shown as a magenta ribbon with stick models of the peptide residues (Y13, F14, I19, I20, and W21). **b, c,**Binding pockets for bosentan (**b**) and IRL2500 (**c**). The bosentan-bound receptor (PDB code: 5XPR) is shown as a thin orange ribbon model. The residues involved in bosentan binding (D154^257^, Q181^3 32^, K182^333^, K273^538^, W3 3 6^6 48^, and R3 4 3 ^6 55^) and D147^250^ are shown as sticks. Bosentan is shown as an orange stick model. IRL2500 and the IRL2500-bound receptor are coloured as in panel (**a**). The residues involved in IRL2500 binding (N158^261^, K182^333^, R343^655^, D147^2 50^, and W336^648^) are shown as sticks.

IRL2500 has distinct chemical moieties as compared with bosentan, because IRL2500 was not developed based on bosentan. To reveal the similarities and differences in their binding modes, we compared the binding modes of IRL2500 and bosentan in detail (Fig. 4b, c). The carboxylate of IRL2500 and the sulfonamide of bosentan are similarly coordinated by the positively charged residue R343^655^, suggesting that this electrostatic interaction is a common feature of the antagonist binding to the ET **B** receptor. In addition, like bosentan, the aromatic moieties of IRL2500 fit within the local hydrophobic pockets in the ET **B** receptor. Overall, IRL2500 has moieties that form similar binding interactions to those of bosentan. However, bosentan lacks the moiety corresponding to the biphenyl group of IRL2500, which deeply penetrates into the receptor core. Thus, IRL2500 fits into the pocket more tightly as compared with bosentan, contributing to its higher affinity.

### IRL2500 function an inverse agonist for ETb

To obtain mechanistic insights into the receptor inactivation by IRL2500, we compared the ET **B** structures bound to ET-1, bosentan, and IRL2500. Previous structural studies showed that ET-1 binding induces the inward moment of the extracellular portion of TM6 including W3 3 6 **^6, 48^**, leading to receptor activation on the intracellular side **^21^**(Fig. 5a). Bosentan binding sterically prevents the inward motion of W3 3 6 **^6, 48^**with its 2-methoxyphenoxy group, and thus functions as an antagonist **^22^**(Fig. 5b). The dimethylphenyl group of IRL2500 superimposes well with the 2-methoxyphenoxy group of bosentan and similarly prevents the inward motion. Moreover, the dimethyl phenyl and biphenyl groups of IRL2500 sandwich the W3 3 6 **^6, 48^**side chain, tightly preventing its inward rotation. These observations suggest that IRL2500 strongly prevents the transition to the active state, as compared with bosentan, thereby possibly working as an inverse agonist that reduces the basal activity.

**Fig.5.**
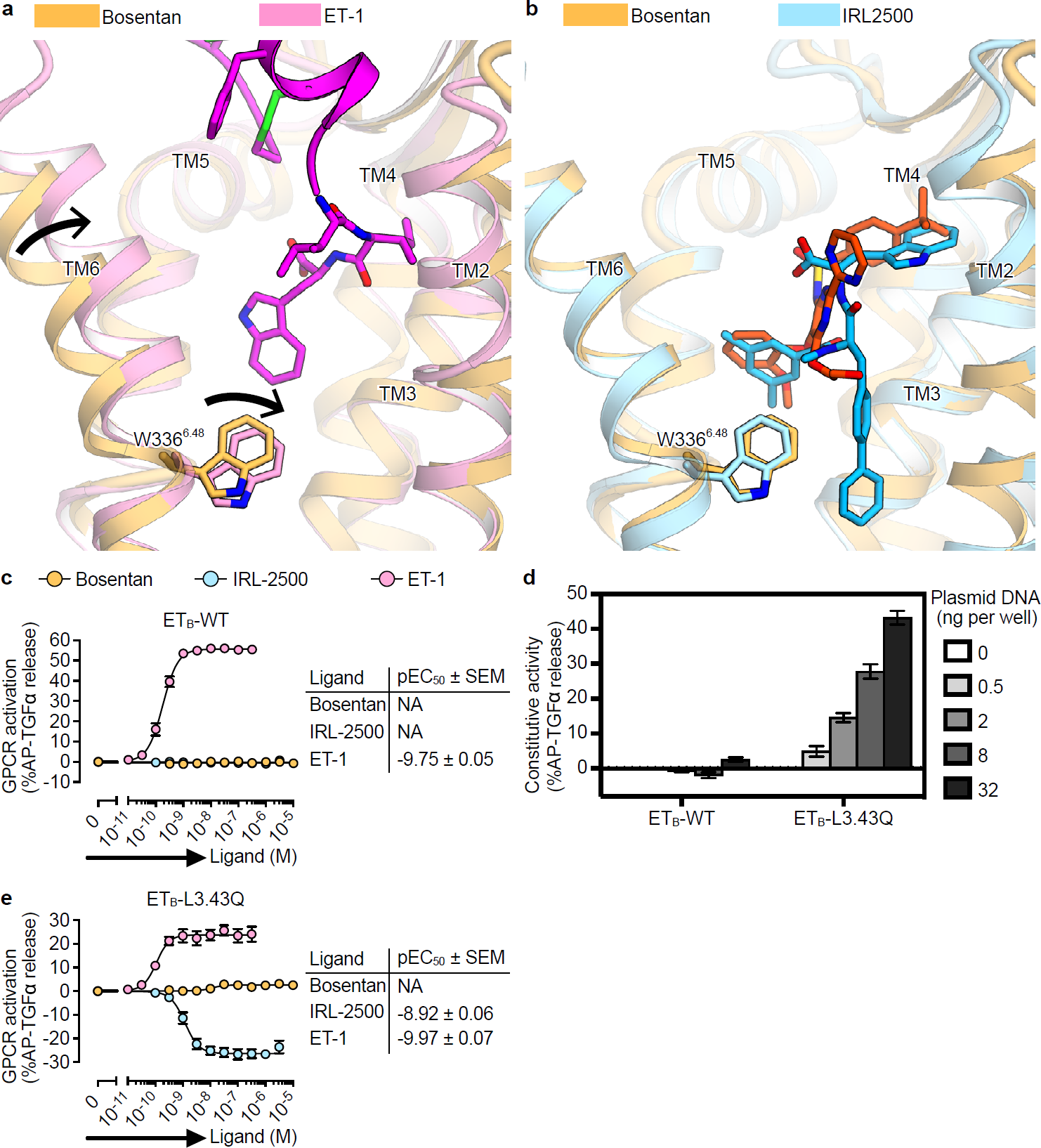
Inverse agonistic activity of IRL2500. **a, b,**Structural changes upon ET-1 and IRL2500 binding, as compared with the bosentan-bound structure, coloured as in Fig. 4. Black arrows indicate the inward movements of TM6 and W3 3 6^6 48^. **c,**Effects of IRL2500 and bosentan on the ET-1-induced AP-TGFa release for the ET **B** receptor. Data are displayed as means ± s.e.m. (standard error of the mean) from three to four independent experiments. **d,**Constitutive activity of ETB. HEK293 cells were transfected with titrated volume of a plasmid encoding the wild-type ET **B** (ET **B**-WT) or the L3.43Q-mutant ET **B** (ET **B**-L3.43Q) and accumulated AP-TGFa release during 24 h after transfection was measured. AP-TGFa release signal in 0 ng receptor plasmid was set as a baseline. Data are displayed as means ± s.e.m. from three independent experiments. **e,**Effects of IRL2500 and bosentan on the ET-1-induced AP-TGFa release for the constitutive active ET **B** receptor (ET **B**-L3.43Q). Data are displayed as means ±s.e.m. from three to four independent experiments.

To investigate the inverse agonist activity of IRL2500 for the ET **B** receptor, we first measured ligand-induced AP-TGFa release responses. The EC **50**value of the agonist ET-1 was 0.11 nM, while IRL2500 and the antagonist bosentan did not change the receptor activation level (Fig. 5c). These data suggested that IRL2500 does not have the inverse agonist activity or that the assay is not sensitive enough to detect inverse agonist activity. Indeed, we observed that the basal activity of the ET **B** receptor was very low in the assay (Fig. 5d) and thus we could not distinguish whether IRL2500 functions as an antagonist or an inverse agonist by this assay.

Therefore, we tried the same assay using a constitutively active mutant of the ET **B** receptor. Constitutively active mutant GPCRs have been employed in pharmacological characterizations of inverse agonists **^30^**, because such mutant GPCRs allow the assay to measure signals in a larger detection window. The substitution of the highly conserved L3.43 to glutamine has been identified as a causative activating mutation in the TSHR **^31^** and CYSLTR2 **^32^**genes, which are related to hyperthyroidism and uveal melanoma, respectively. Therefore, we transferred the L3.43Q mutation into ET **B** (ET **B**-L192 **^3.43^**Q) and examined its constitutive activity. We found that ET **B**-L3.43Q induced spontaneous AP-TGFa release in a plasmid volume-dependent manner (Fig. 5d), indicating that L3.43Q works as a constitutive active mutation in the ET **B** receptor. We evaluated the dose response effects of bosentan and IRL2500, using the constitutive active mutant ET **B**-L3.43Q (Fig. 5e). Again, the antagonist bosentan did not change the receptor activation from the baseline level, whereas IRL2500 reduced the basal activity (EC **so**=1.2 nM). These data indicate that IRL2500 works as a potent inverse agonist for the ET **B**receptor, consistent with the structural observations. The biphenyl group of IRL2500 prevents the inward motion of W336 **^6.48^**and stabilizes the inactive conformation, and thus IRL2500 functions as an inverse agonist.

## Discussion

We have determined the crystal structure of the ETB receptor in complex with the peptide compound IRL2500, and thus elucidated the detailed receptor interactions and the structural basis for its ETB selectivity. Although IRL2500 is designed to mimic the partial structure of ET-1, the binding mode is quite different. Moreover, using the constitutively active mutant that we established in the current study, we first revealed that IRL2500 functions as a potent inverse agonist for the ET **B**receptor, and provided the structural basis for the inverse agonistic mechanism.

Small-molecule ETR antagonists have been developed over the years; however, most ETR antagonists have been designed based on bosentan. Thus, the presently available ET agents are chemically very similar. IRL2500 was developed based on ET-1 and has totally distinct chemical moieties, as compared with bosentan. However, the comparison of the IRL2500 and bosentan binding modes revealed the unexpected similarity in their binding interactions. This observation suggests that the chargecomplementary interactions in the center of the pocket form the core of the receptor-antagonist interactions, and the other aromatic moieties fit the local hydrophobic pocket. The ligand binding pocket in the inactive ET **B**structures is larger than those in other GPCR structures, and thus aromatic moieties may be necessary to fit well within the pocket.

We revealed that the biphenyl group of IRL2500 penetrates deeply into the receptor core proximal to D147 **^2, 50^**, preventing the inward motion of W336 **^6.48^**in TM6, and thus IRL2500 functions as an inverse agonist (Fig. 6a). This D2.50 constitutes a sodium binding site adjacent to the orthosteric site, which is highly conserved among the class A GPCRs^33^. Sodium has negative allosteric effects on ligand binding in most class A GPCRs, by stabilizing the inactive conformations. Therefore, this sodium binding site is the hot spot for the design of allosteric modulators and inverse agonists to fix receptors in the inactive conformations. In the BLT1 structure bound to the inverse agonist BIIL260, the benzamidine group of BIIL260 directly hydrogen bonds with D2.50, stabilizing the inactive conformation instead of the sodium^34^ (Fig. 6b). The biphenyl group of IRL2500 superimposes well with the benzamidine group of BIIL260 (Fig. 6c). Although the biphenyl group of IRL2500 does not form any hydrogen-bonding interactions with the receptor, it prevents the conformational change around the D2.50 in a similar manner to the benzamidine moiety of BIIL260. For the design of effective inverse agonists, the biphenyl moiety would be also useful as a modulation part along with another moiety that exerts specific and tight binding to the orthosteric site, as well as a benzamidine group.

**Fig.6.**
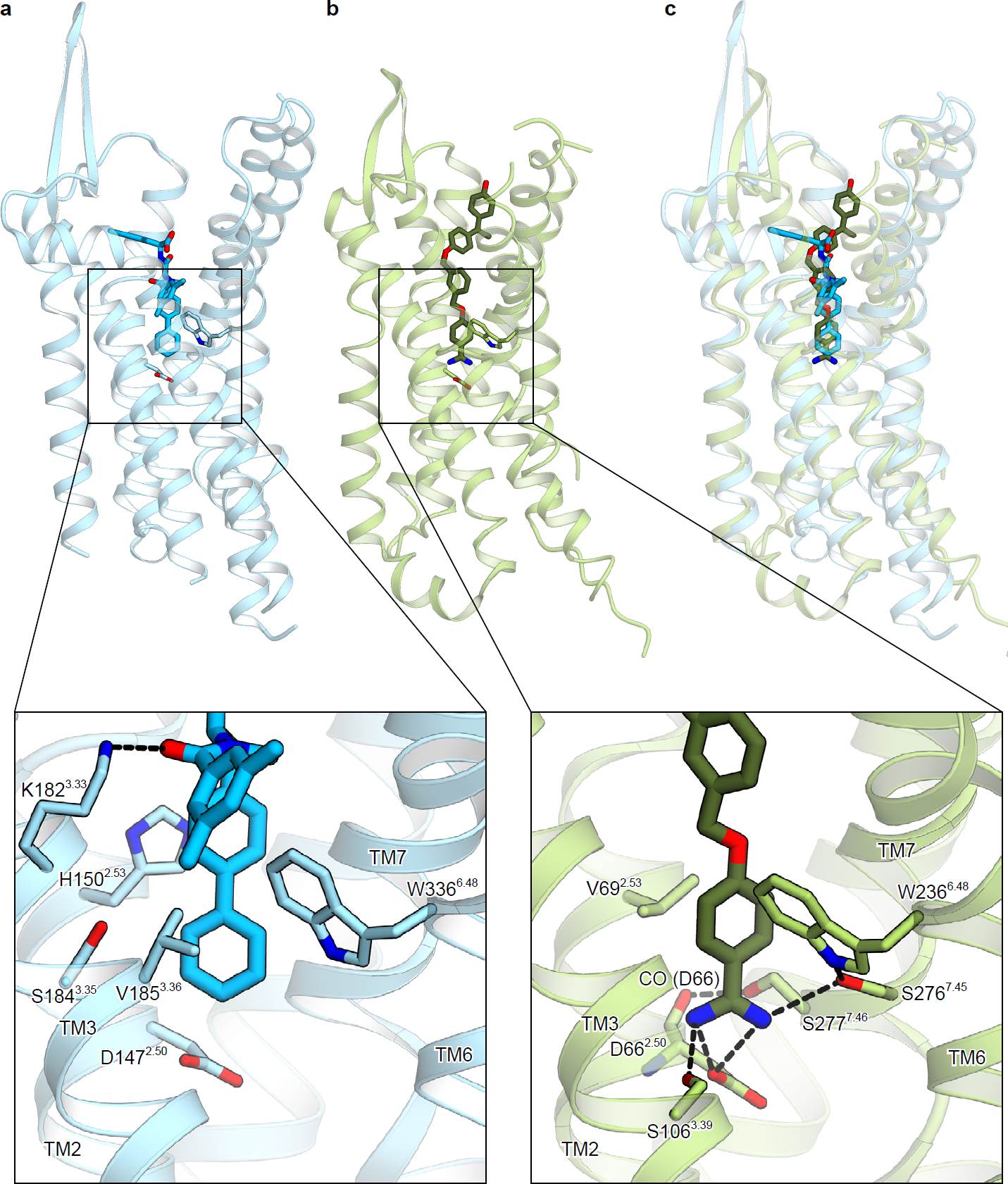
Structural comparison of ETb and BLT1 in complex with inverse agonists. **a, b,**The overall structures of the IRL2500-bound ET **B** receptor (**a**) and the BIIL260-bound BLT1 receptor (PDB code: 5X33) (**b**). The ET **B** and BLT1 receptors are shown as sky blue and light green ribbons, respectively. The inverse agonists IRL2500 and BIIL260 are shown as blue and dark green sticks, respectively. The lower panels show the binding interactions around the sodium binding site. The residues involved in ligand binding are represented with sticks. Hydrogen bonds are indicated by black dashed lines. In the IRL2500-bound structure, the biphenyl group of IRL2500 forms van der Waals interactions with the receptor, and does not form any hydrogen-binding interactions. In the BIIL260 structure, BIIL260 forms hydrogen bonds with D66 ^250^, S106^339^, and S276^745^. **c,**Superimposition of the ETB and BLT1 structures in complex with the inverse agonists.

### Materials and methods

#### Expression and purification

The haemagglutinin signal peptide, followed by the Flag epitope tag (DYKDDDDK) and a nine-amino-acid linker, was added to the N-terminus of the receptor, and a tobacco etch virus (TEV) protease recognition sequence was introduced between G57 and L66, to remove the disordered N-terminus during the purification process. The C-terminus was truncated after S407, and three cysteine residues were mutated to alanine (C396A, C400A, and C405A) to avoid heterogeneous palmitoylation. To improve crystallogenesis, we introduced four thermostabilizing mutations () and inserted minimal T4 lysozyme **^25^**into intracellular loop 3, between L303 **^5.68^**and L311 **^6.23^**(ET **B**-Y4-mT4L **^22^**).

The thermostabilized construct ET **B**-Y4-mT4L was subcloned into a modified pFastBac vector, with the resulting construct encoding a TEV cleavage site followed by a GFP-His **^10^**tag at the C-terminus. The recombinant baculovirus was prepared using the Bac-to-Bac baculovirus expression system (Invitrogen). Sf9 insect cells were infected with the virus at a cell density of 4.0 × 10 **^6^**cells per millilitre in Sf900 II medium, and grown for 48 h at 27 °C. The harvested cells were disrupted by sonication, in buffer containing 20 mM Tris-HCl, pH 7.5, and 20% glycerol. The crude membrane fraction was collected by ultracentrifugation at 180,000g for 1 h. The membrane fraction was solubilized in buffer, containing 20 mM Tris-HCl, pH 7.5, 200 mM NaCl, 1% DDM, 0.2% cholesterol hemisuccinate, 10 μM IRL2500, and 2 mg ml **^-1^**iodoacetamide, for 1 h at 4 °C. The supernatant was separated from the insoluble material by ultracentrifugation at 180,000g for 20 min, and incubated with TALON resin (Clontech) for 30 min. The resin was washed with ten column volumes of buffer, containing 20 mM Tris-HCl, pH 7.5, 500 mM NaCl, 0.1% LMNG, 0.01% CHS, 10 pM IRL2500, and 15 mM imidazole. The receptor was eluted in buffer, containing 20 mM Tris-HCl, pH 7.5, 500 mM NaCl, 0.01% LMNG, 0.001% CHS, 10 μM IRL2500, and 200 mM imidazole. The eluate was treated with TEV protease and dialysed against buffer (20 mM Tris-HCl, pH 7.5, 500 mM NaCl, and 10 μM IRL2500). The cleaved GFP-His **10**tag and the TEV protease were removed with Co **^2^+**-NTA resin. The receptor was concentrated and loaded onto a Superdex200 10/300 Increase size-exclusion column, equilibrated in buffer containing 20 mM Tris-HCl, pH 7.5, 150 mM NaCl, 0.01% LMNG, 0.001% CHS, and 10 μM IRL2500. Peak fractions were pooled, concentrated to 40 mg ml **^-1^**using a centrifugal filter device (Millipore 50kDaMW cutoff), and frozen until crystallization. During the concentration, IRL2500 was added to a final concentration of 100 pM.

#### Crystallization

The purified receptor was reconstituted into molten lipid (monoolein and cholesterol 10:1 by mass) at a weight ratio of 1:1.5 (protein:lipid). The protein-laden mesophase was dispensed into 96-well glass plates in 30 nl drops and overlaid with 800 nl precipitant solution by a Gryphon LCP robot (Art Robbins Instruments) **^26^**. Crystals of ET **B**-Y4-mT4L bound to IRL2500 were grown at 20°C in precipitant conditions containing 30% PEG300, 100 mM Bis-tris, pH 7.5, 150 mM sodium phosphate monobasic, and 10 mM TCEP hydrochloride. The crystals were harvested directly from the LCP using micromounts (MiTeGen) or LithoLoops (Protein Wave) and frozen in liquid nitrogen, without adding any extra cryoprotectant.

#### Data collection and structure determination

X-ray diffraction data were collected at the SPring-8 beamline BL32XU, with 10 × 15 ^2^ (width × height) micro-focused beams and an EIGER × 9M detector (Dectris). Various wedge data sets (10°) per crystal were mainly collected with the ZOO system, an automatic data-collection system developed at SPring-8 (K.Y., G.U., K.H., M.Y., and K.H., submitted). The loop-harvested microcrystals were identified by raster scanning and subsequently analyzed by SHIKA^35^. Each data set was indexed and integrated with XDS^36^, and the datasets were hierarchically clustered by using the correlation coefficients of the intensities between datasets. After the rejection of outliers, 58 data sets were finally merged with XSCALE^36^. The IRL2500-bound structure was determined by molecular replacement with PHASER^37^, using the K8794-bound ET **B** structure (PDB code: 5X93). Subsequently, the model was rebuilt and refined using COOT^38^ and PHENIX^39^, respectively. The final model of IRL2500-bound ET **B**-Y4-T4L contained residues 91-207, 214-303, and 311-403 of ET **B**, 1-14 and 22-117 of mT4L, IRL2500, 6 monoolein molecules, two phosphoric acids, and 41 water molecules. The model quality was assessed by MolProbity^40^. Figures were prepared using CueMol (http://www.cuemol.org/ja/)

#### TGFa shedding assay

The TGFa shedding assay, which measures the activation of Gq and G12 signaling^24^, was performed as described previously^22^. Briefly, a plasmid encoding an ET **B** construct with an internal FLAG epitope tag or an ET **A** construct was transfected, together with a plasmid encoding alkaline phosphatase (AP)-tagged TGFα (AP-TGFα), into HEK293A cells by using a polyethylenimine (PEI) transfection reagent (1 gg ETR plasmid, 2.5 gg AP-TGFa plasmid, and 25 μl of 1 mg/ml PEI solution per 10-cm culture dish). After a one day culture, the transfected cells were harvested by trypsinization, washed, and resuspended in 30 ml of Hank’s Balanced Salt Solution (HBSS) containing 5 mM HEPES (pH 7.4). The cell suspension was seeded in a 96 well plate (cell plate) at a volume of 80 μl per well and incubated for 30 min in a CO **2**incubator. For the measurement of antagonist activity, IRL2500 was diluted in 0.01% bovine serum albumin (BSA) and HEPES-containing HBSS (assay buffer) and added to the cell plate at a volume of 10 gl per well. After 5 min, ET-1, at a final concentration of 0.2 nM, was added to the cell plate at a volume of 10 μl per well. For the measurement of agonistic activity, after adding 10 gl of the assay buffer, serially diluted ET-1 was mixed with the cells at a volume of 10 gl per well. After a 1 h incubation in the CO **2**incubator, aliquots of the conditioned media (80 μl) were transferred to an empty 96-well plate (conditioned media (CM) plate). Similarly, for the measurement of inverse agonist activity, the cells were mixed with 10 gl of the assay buffer, followed by the addition of serially diluted IRL2500, and incubated for 4 h before the transfer of the conditioned media. The AP reaction solution (10 mM p-nitrophenylphosphate (p-NPP), 120 mM Tris-HCl (pH 9.5), 40 mM NaCl, and 10 mM MgCh) was dispensed into the cell plates and the CM plates (80 μl per well). The absorbance at 405 nm (Abs **405**) of the plates was measured, using a microplate reader (SpectraMax 340 PC384, Molecular Devices), before and after a 1 h incubation at room temperature. AP-TGFα release was calculated as described previously **^22^**. The AP-TGFa release signals were fitted to a four-parameter sigmoidal concentration-response curve, using the Prism 7 software (GraphPad Prism), and the pEC **50**(equal to-Log **i0**EC **50**) and E **max**values were obtained.

To measure the constitutive activity in a plasmid volume-dependent manner, HEK293 cells were seeded in a 96-well plate at a concentration of 4 × 10 **^5^**cells per ml in Opti-MEM I Reduced Serum Media (Thermo Fisher Scientific), in a volume of 80 gl per well. A transfection mixture was prepared by mixing the PEI transfection reagent (0.2 gl per well) and plasmids (20 ng AP-TGFα plasmid, titrated ETR plasmid, and an empty vector to balance the total plasmid volume) in Opti-MEM I Reduced Serum Media (20 gl). The mixture was added to the cells, which were then incubated for 24 h before the transfer of the conditioned media. After adding the AP reaction solution, the absorbances of the cells and the CM plates were measured at 20 min intervals. The AP-TGFa release signals were calculated as described above, and the signal in the mock-transfected conditions was set at the baseline.

## Acknowledgements

The diffraction experiments were performed at SPring-8 BL32XU (proposal 2017A2527). We thank the beamline staff at BL32XU of SPring-8 (Sayo, Japan) for technical assistance during data collection. We also thank Kouki Kawakami, Takeaki Shibata and Ayumi Inoue (Tohoku University, Japan) for technical assistance in the characterization of the L3.43Q-mutant ET **B** receptor. This work was supported by grants from the Platform for Drug Discovery, Informatics and Structural Life Science by the Ministry of Education, Culture, Sports, Science and Technology (MEXT), JSPS KAKENHI grants 16H06294 (O.N.), 17J30010 (W.S.), 30809421 (W.S.), 17K08264 (A.I.), and the Japan Agency for Medical Research and Development (AMED) grants: the PRIME JP17gm5910013 (A.I.) and the LEAP JP17gm0010004 (A.I. and J.A.), and the National Institute of Biomedical Innovation.

## Author contributions

C.N. expressed, purified, and crystallized the IRL2500-bound ET **B** receptor, collected data, and refined the structures. W.S. designed all of the experiments, initially crystallized the receptor, and refined the structure. A.I., F.M.N.K., and J.A. performed and oversaw the cell-based assays. The manuscript was prepared by C.N., W.S., A.I., and O.N. W.S. and O.N. supervised the research. Coordinates and structure factors have been deposited in the Protein Data Bank, under the accession number XXXX for the IRL2500-bound structure. The X-ray diffraction images are also available at SBGrid Data Bank (https://data.sbgrid.org/), under the ID YYYY.

